# TGFβ determines epithelial tissue spacing by regulating mesenchymal condensation

**DOI:** 10.64898/2026.03.16.712215

**Authors:** Chan Jin Park, Pengfei Zhang, Carolina Trenado-Yuste, Celeste M. Nelson

## Abstract

The branched structure of the vertebrate lung provides a high surface area-to-volume ratio, which increases the efficiency of diffusion-driven gas exchange. Generating this structure requires that the epithelial branches avoid contacting each other as they elongate during development. Previous studies have suggested that the spacing between neighboring branches is intrinsic to growth of the epithelium, but the underlying physical mechanisms remain elusive. Here, we used the embryonic chicken lung as a model system and found that branch spacing is regulated primarily by signaling to the mesenchyme through transforming growth factor-beta (TGFβ). Although proliferation decreases in epithelial cells that are located in close proximity to an adjacent branch, these patterns surprisingly emerge after regular branch spacing has been established. Instead, we find that TGFβ promotes the directed migration of mesenchymal cells, which form a condensation that physically displaces the adjacent epithelium and tunes branch spacing. Continuous disruption of TGFβ signaling prevents mesenchymal condensation and eventually results in contact between adjacent branches. These data suggest that the spacing between epithelial branches results from mesenchymal cell dynamics rather than from epithelial-intrinsic self-avoidance.

## Introduction

Tissue shape is closely related to tissue function. A paradigmatic example is the branched epithelial tube, which is commonly found in organs that promote the exchange of material, including the lung, mammary gland, kidney, and salivary gland.^1–3^ Branched epithelia maximize surface area within a confined volume and enable the efficient delivery of gases, salts, and proteins.^4^ These epithelial trees are constructed from simpler epithelial precursors through a process known as branching morphogenesis.^1,4,5^ In the mammalian and avian lungs, the recursive process of branch initiation, elongation, and termination leads to the formation of ramified tree-like structures.^4,6,7^

One often overlooked aspect of lung development is that, despite being densely packed, the epithelial branches remain separated from each other during the elongation phase of branching morphogenesis.^4^ Although this self-avoidance is critical for maximizing the area of the gas-exchange surface, it is unclear how epithelial branches within a developing lung sense and avoid colliding with each other. In the mammary gland, transforming growth factor (TGF)-β produced by the epithelium terminates the growth of adjacent branches.^8^ In the kidney, biochemical signaling downstream of the TGFβ-family member, bone morphogenetic protein (BMP)-7, inhibits epithelial proliferation in neighboring branches.^9^ In the salivary gland, a decrease in proliferating epithelial cells in branches closely apposed to each other is also observed, but the underlying molecular mechanism remains elusive.^10^ Based on this experimental evidence, epithelial spacing in branched organs is generally assumed to result from interactions between adjacent branches that inhibit epithelial proliferation as they approach each other, although different organs may employ distinct molecular pathways.^8–10^

Here, we used the embryonic chicken as a model system to study how epithelial branches sense and avoid colliding with each other in the developing lung. We took advantage of the fact that the spacing between branches is uniform in the embryonic chicken lung, even at early stages of development. Using immunofluorescence analysis, surgical manipulations, pharmacological inhibitors, and timelapse imaging, we found that epithelial proliferation in neighboring branches is inhibited by signaling through TGFβ but not BMP. However, regular branch spacing is established even before epithelial proliferation decreases, implying an additional mechanism to prevent epithelial contact. Consistently, we found that mesenchymal cells migrate toward and form stiff condensations around focal sources of TGFβ. These mesenchymal condensations physically displace the adjacent epithelium, acting as a determinant of branch spacing, an effect that we also observed in the embryonic mouse lung and salivary gland. Conversely, continuously disrupting TGFβ signaling eventually leads to thinning of the mesenchyme and contact between adjacent branches. These data suggest that TGFβ-mediated signaling affects both epithelial and mesenchymal cells and is essential for branches within the developing chicken lung to avoid colliding with each other, thus allowing the organ to achieve a high surface area-to-volume ratio.

## Materials and Methods

### Experimental model

All procedures involving animals were approved by Princeton University’s Institutional Animal Care and Use Committee. Fertilized White Leghorn chicken eggs (Department of Animal Science at the University of Connecticut) and transgenic cytoplasmic GFP chicken eggs (Susan Chapman, Clemson University) were incubated at 37.5°C in a humidified atmosphere until the desired stages. Lungs were surgically removed from embryos between embryonic days 5 (*E*5) and 8 (*E*8), which corresponds to Hamburger-Hamilton stages (HH) 25–34.^11^ CD1 mice were maintained under standard laboratory conditions. Lungs, kidneys, and salivary glands were surgically removed from *E*12.5, *E*13.5, and *E*14.5 mouse embryos, respectively. Noon of the day on which a vaginal plug was detected was considered as embryonic day 0.5 (*E*0.5).

### Ex-vivo culture and live-imaging of embryonic organs

Microdissected lung, kidney, and salivary gland explants were cultured *ex vivo* following established protocols.^12^ Briefly, tissue explants were cultured at the air-liquid interface on a porous membrane floating on DMEM/F12 medium (without HEPES; Gibco, 21041025) supplemented with 5% fetal bovine serum (FBS; R&D Systems, S11150H) and antibiotics (50 U/ml of penicillin and streptomycin; Thermo Fisher Scientific, 15070063). Tissue explants were cultured in an incubator maintained at 37°C and 5% CO_2_. For studies using exogenous growth factors, fibroblast growth factor-10 (FGF10; R&D Systems, 345-FG) was first dissolved in sterile phosphate-buffered saline (PBS) containing 0.1% bovine serum albumin (BSA; Sigma-Aldrich, A7906). Dissolved FGF10 was then added to the culture medium at a final concentration of 1 µg/ml. PBS containing 0.1% BSA was used as a vehicle control. For studies using inhibitors, RepSox (Sigma-Aldrich, R0158) or dorsomorphin (Sigma-Aldrich, P5499) were first dissolved in DMSO and then added to the culture media. Aphidicolin (Sigma-Aldrich, 75351), PF-573228 (Selleck Chemicals, S2013), and para-amino-blebbistatin (Cayman Chemical, 22699) were dissolved in dimethylformamide (DMF). For continuous inhibition experiments, the media was replaced every 24 hours. Either DMSO or DMF were used as vehicle controls. For live-imaging, microdissected lungs were placed on a glass-bottom plate and imaged on a multiphoton microscope (Nikon AX R MP) equipped with a tabletop incubator.

### Preparation of TGFβ1-containing beads

Recombinant human transforming growth factor-beta1 (TGFβ1; HEK293 derived; PeproTech, 100-21) was reconstituted in sterile 4-mM HCl containing 0.1% BSA. Agarose beads (75-150-µm diameter; Bio-Rad, 1537301) were rinsed with PBS for 30 minutes at room temperature. The beads were then incubated overnight at 4°C in either reconstitution buffer alone as a control or in solutions containing 20- or 100-µg/ml TGFβ1.

### Immunofluorescence analysis

For 5-ethynyl-2’deoxyuridine (EdU)-incorporation experiments, lung explants were pulsed for 30 minutes prior to fixation unless stated otherwise. Fixed samples were stained following the instructions of the Click-iT EdU Alexa Fluor 555 Kit (Invitrogen, C10638) prior to incubating with primary antibodies.

Samples were fixed with 4% paraformaldehyde for 30 minutes, rinsed with Tris-buffered saline (TBS), and washed three times for 15 minutes each with 0.5% Triton X-100 in TBS (TBST). Samples were blocked with BlockAid^™^ Blocking Solution (Invitrogen, B10710) and TBST mixture (50% v/v) for 1 hour. Samples were then incubated with primary antibodies against E-cadherin (1:100; BD Biosciences, 610182), pSMAD1/5/9 (1:100; Invitrogen, PA5-64711), pan-laminin (1:100; Invitrogen, PA1-16730), perlecan (1:100; Invitrogen, MA1-06821), or pSMAD2 (1:100; Cell Signaling, 18338). Samples were washed four times for 15 minutes each with TBST, blocked again for 1 hour, and then incubated with DAPI (3:500; Invitrogen, D1306), Alexa Fluor 488 phalloidin (Thermo Fisher Scientific, A12379), and Alexa Fluor-conjugated secondary antibodies (3:500; Invitrogen, A32766 or A32795).

### Immunoblotting analysis

Samples were lysed in RIPA lysis buffer supplemented with cOmplete™ protease inhibitor cocktail (Roche, 11836170001) and Pierce™ phosphatase inhibitor cocktail (Thermo Fisher Scientific, A32957). Protein concentrations were measured using a Pierce™ BCA protein assay kit (Thermo Fisher Scientific, 23227). Samples were then mixed with NuPAGE™ LDS sample buffer (Thermo Fisher Scientific, NP0007) and NuPAGE™ sample reducing agent (Thermo Fisher Scientific, NP0004), boiled at 95°C for 10 minutes, resolved by SDS-PAGE, and transferred to nitrocellulose membranes. Membranes were blocked in 5% BSA in 0.1% Tween-20 in TBS overnight at 4°C and then incubated in blocking buffer containing antibodies against pSMAD2 (1:1000; Thermo Fisher Scientific, 44-244G), pERK1/2 (1:1000; Cell Signaling, 4370), or GAPDH (1:1000; Cell Signaling, 3683) for 2 hours at room temperature. Bands were detected using horseradish-peroxidase-conjugated secondary antibody (1:1000, Cell Signaling, 7074) and Amersham ECL Western Blotting Reagent (GE Healthcare) as a chemiluminescent substrate. Densitometry analysis was performed using ImageJ.

### Fluorescence in situ hybridization

Samples were fixed with 4% paraformaldehyde for 24 hours at 4°C, then sequentially washed in RNase-free water, followed by 20% and 30% sucrose solution in RNase-free water. Samples were then embedded in OCT (Tissue Tek) using dry ice and sectioned at a thickness of 10 μm with a cryostat (Leica, CM3050S). Fluorescence in situ hybridization was performed using the standard RNAScope Multiplex Fluorescent V2 ASSAY protocol (ACD). Probes used were for *Gallus gallus Fgf10* (1596441)*, Fgfr2* (1106541)*, Tgfb1* (534751)*, Tgfb2* (1055431)*, Tgfb3* (1596461)*, Tgfbr1* (1596431)*, and Tgfbr2* (1596451).

### Spatial transcriptomics

Spatial transcriptomic mapping was performed using a modified microfluidic-enabled, deterministic barcoding-based (DBiT-seq) workflow.^13–15^ Dissected lungs were embedded in OCT, sectioned at a thickness of 12 μm with a cryostat, and transferred onto poly-L-lysine-coated glass slides (Electron Microscopy Sciences, 63478-AS). The sections were spatially barcoded through two rounds of barcode application using microfluidic devices with a channel width of 15 μm. After in situ ligation, barcoded tissue sections were digested and cDNA was purified before being sequenced.

### Stiffness measurements

Snap-frozen lung explants were embedded in OCT using dry ice and sectioned at a thickness of 12 μm with a cryostat. Cryosections were transferred onto microscope slides (Fisher Scientific, Fisherbrand™ Superfrost™ Plus), which were then mounted to the bottom of a PBS-filled 1-well plate and loaded into a Pavone Nanoindenter (Optics 11 life). Measurements were performed using a probe with tip radius of 9.5 μm and stiffness of 0.017 N/m, assuming a Poisson’s ratio of 0.5. Peak-load poking mode with max load of 0.1 μN was used to measure tissue stiffness. For matrix scans, the *x*- and *y*-spacing between adjacent points was set to 40 μm.

### Agent-based modeling

We used a two-dimensional (2D) agent-based model to simulate the collective behavior of mesenchymal cells and their interactions with an epithelial branch.^16,17^ Cells were represented as point particles defined by their centers of mass. The simulation domain was a rectangular box of size 20×20 with reflecting boundary conditions. A source of activating morphogen was placed at the center of the domain, and the source-adjacent region was defined as a circular region around the source with a radius of *R*.

The positions of mesenchymal cells were updated at every timestep, where the trajectory of a mesenchymal cell *i* was determined according to a Langevin-like equation:

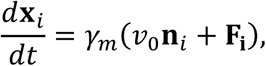

where **x**_*i*_ is the position of the cell, *γ*_*m*_ is the motility coefficient of a mesenchymal cell, *v*_0_ is the self-propulsion speed, **n**_*i*_ = (cos *θ*_*i*_, sin *θ*_*i*_) is the polarity vector, and 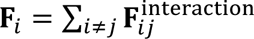 is the net interaction force applied on the cell. In the model, cells experience short-range repulsive interactions to prevent overlap, where the pairwise force between cells *i* and *j* whose intercellular distance *r*_*ij*_ is smaller than the cutoff distance *d*_*c*_ = 0.6 was defined as:

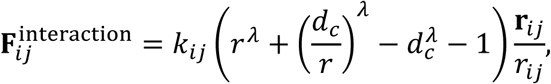

where *k*_*ij*_ is the repulsion strength and *λ* = 2. Cells with intercellular distance larger than *d*_*c*_ were assumed to not interact with each other.

To model directed migration, the polarity vector of cells within the source-adjacent region was adjusted towards the source as a function of distance from the source *d*_*i*_:

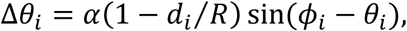

where *α* is the chemotactic strength and *ϕ*_*i*_ is the angle between the source and the cell. This adjustment implies that the direction of propulsion of the cells is strongly oriented near the source.

To model enhanced proliferation, new cells were added within the source-adjacent region. The number of added cells was determined by the product of the current number of cells within the source-adjacent region and the proliferation coefficient *p* (0 < *p* < 1).

Each epithelial branch was modeled as a deformable circular chain consisting of epithelial cells. The equation of motion for an epithelial cell *i* was described as:

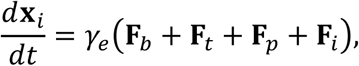

where *γ*_*e*_ is the motility coefficient of an epithelial cell and **F**_*b*_, **F**_*t*_, **F**_*p*_ are the bending, tension, and pressure forces, respectively. The bending force was incorporated to align the cells and obtain a smooth epithelial tissue outline, which was defined as:

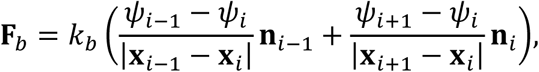

where *k*_*b*_ is the bending elasticity coefficient, *ψ*_*i*_ is the angle between vectors **x**_*i*_ − **x**_*i*−1_, **x**_*i*+1_ − **x**_*i*_, and **n**_*i*_ is the unit vector normal to **x**_*i*+1_ − **x**_*i*_. The tension force was incorporated to maintain cohesion, which was defined as:

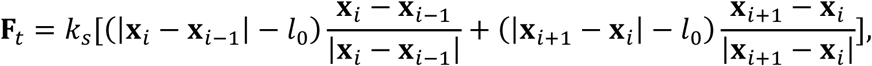

where *k*_*s*_ is the tension coefficient and *l*_0_ is the resting length, which is the length set at initialization. The luminal pressure within the branch was described with the pressure force, which was defined as:

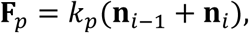

where *k*_*p*_ represents its magnitude.

## Results

### Epithelial proliferation decreases while mesenchymal proliferation is maintained adjacent to neighboring branches

In the embryonic chicken lung, approximately 20 secondary bronchi emerge from each primary bronchus between embryonic days 5 and 8 (*E*5-8) (**Fig. 1A, B**).^7^ During this same period of time, secondary bronchi elongate into the surrounding mesenchyme (**Fig. 1C, D**). We found that from *E*6 to *E*8, the branches are oriented parallel to each other and separated by a nearly constant distance of ∼25 μm (**Fig. 1C, E, F**). This average spacing between secondary bronchi is comparable to the critical distance at which epithelial branches are thought to sense each other in several other organs, including the mammalian kidney and salivary gland.^9,10,18,19^

**Fig. 1.**
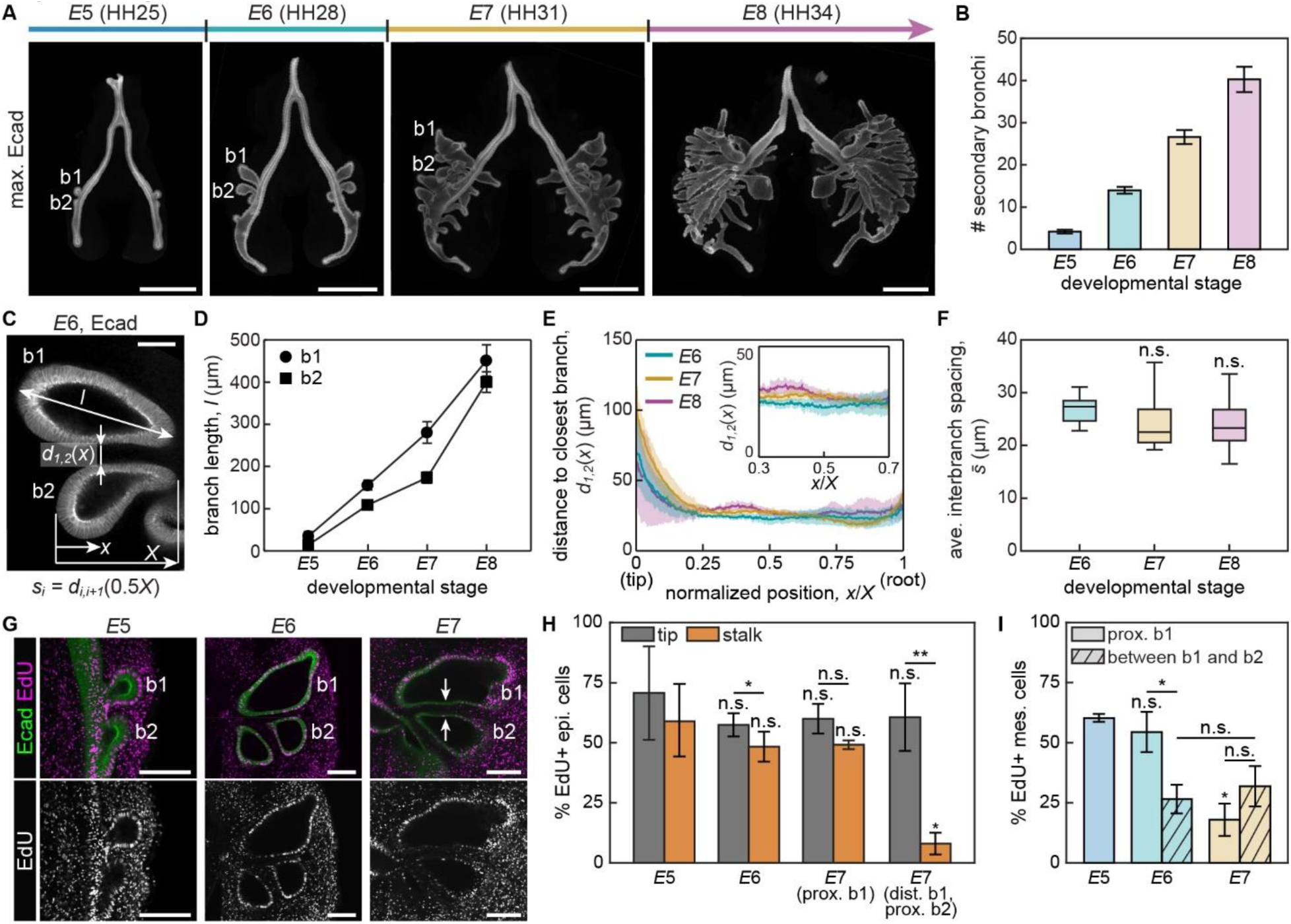
The spacing between branches remains constant while the spatial patterns of epithelial and mesenchymal proliferation change significantly over developmental time in the embryonic chicken lung. A. Maximum-intensity projection of immunofluorescence for E-cadherin in embryonic chicken lungs at different stages of development. b1 and b2 denote the first and second secondary bronchi, respectively. Scale bars, 500 µm. B. Graph showing the number of secondary bronchi as a function of developmental time. Shown are the average and s.d. of at least 4 lungs per timepoint. C. Confocal section of immunofluorescence for E-cadherin from a *E*6 chicken lung depicting variables related to branch geometry. Scale bar, 50 µm. D. Graph showing the lengths of b1 and b2 as a function of developmental time. Shown are the average and s.d. of 3 lungs per timepoint. E. Graph showing the distance (*d_1,2_*) between b1 and b2 over developmental time. Inset shows the stalk region where *d_1,2_* is maintained nearly constant regardless of developmental stage. *x*-axis starts from the tip of b2 and the distance from this tip to the root of the branch is denoted as *X*. Shown are the average and s.d. of 3 lungs per timepoint. F. Graph showing the average spacing between branches over developmental time. Shown are the average and s.d. of 3 lungs per timepoint. n.s. stands for not significant. G. Florescence images of EdU incorporation (magenta) in embryonic chicken lungs at different stages of development, counterstained for E-cadherin (green). White arrows denote closely apposed branch stalks with decreased proliferation. Scale bars, 100 µm. H. Graph showing the percentage of EdU-positive cells in the tips and stalks of the epithelium. prox. and dist. denote the proximal and distal sides of each branch, respectively. Shown are the average and s.d. of 3 lungs per timepoint. Statistical analyses are performed in comparison to *E*5 unless denoted otherwise. *, *p*<0.05. **, *p*<0.005. I. Graph showing the percentage of EdU-positive cells in the mesenchyme. Solid and hatched bars denote percentage of EdU-positive cells proximal to b1 and in between b1 and b2. Since b2 starts to emerge at *E*5, the percentage of EdU-positive cells in between b1 and b2 is measured starting on *E*6. Statistical analyses are performed in comparison to *E*5 unless denoted otherwise.

A constant distance between branches could be maintained if epithelial growth were restricted to the direction of branch extension. To determine whether the parallel spacing of secondary bronchi correlates with patterns of epithelial proliferation, we conducted EdU-incorporation assays in freshly explanted lungs (**Fig. 1G**). This analysis revealed that each secondary bronchus is highly proliferative at its tip, but the epithelial cells within closely apposed stalks become quiescent over developmental time. Specifically, we found that EdU-positive cells are evenly distributed throughout the first (b1) and second (b2) secondary bronchi at *E*5 and *E*6 (**Fig. 1G, H**). At *E*7, however, the percentage of EdU-positive cells drastically decreases in the epithelium located in the distal stalk of b1 and proximal stalk of b2, which are positioned adjacent to each other. In contrast, the percentage of EdU-positive cells remains high in the epithelium located in the proximal stalk of b1, which lacks a neighboring branch. We found a similar pattern of proliferation within the secondary bronchi of lungs isolated at *E*6 and then cultured for 24 hours, regardless of the duration of the EdU pulse (**Fig. S1A, B**).

We also analyzed proliferation in the mesenchyme that surrounds the epithelial branches (**Fig. 1I**). At *E*5 and *E*6, the percentage of EdU-positive mesenchymal cells was similar to the percentage of EdU-positive epithelial cells in the tips of the branches. By *E*7, however, we observed a change in the pattern of proliferation. Specifically, the percentage of EdU-positive cells in the mesenchyme decreased, but the percentage between b1 and b2 was still significantly higher than (more than double) that of the adjacent epithelium. We noted a similar pattern of proliferation within the mesenchyme of lungs isolated at *E*6 and cultured for 24 hours (**Fig. S1C**). These findings suggest that the spacing between branches in the embryonic chicken lung is associated with differential patterns of proliferation in the epithelium and mesenchyme.

### Surgically increasing the spacing between branches prevents the changes in epithelial and mesenchymal proliferation

The above-described patterns of epithelial proliferation, especially the contrast between the highly proliferative proximal stalk of b1 (which lacks a neighboring branch) and the relatively nonproliferative distal stalk of b1 (which is closely apposed to another branch), are highly suggestive of branch-branch communication. To test whether the local decrease in epithelial proliferation results from interactions between adjacent branches, we surgically removed either b1 or b2 from *E*6 lung explants and monitored the pattern of EdU incorporation in the neighboring epithelium (**Fig. 2A**). Because each lung has two sets of secondary bronchi, we only removed a branch from one side of the lung and used the contralateral side as an internal control. After surgery, we cultured each explant for 24 hours before conducting EdU-incorporation assays. We found that the density of EdU-positive cells is significantly higher in the stalks when the neighboring branch is surgically removed (**Fig. 2B**). In the contralateral controls, however, we observed a reduction in proliferation in stalks adjacent to neighboring branches (**Fig. 2B**), identical to the patterns observed in intact lungs (**Fig. 1G**).

**Fig. 2.**
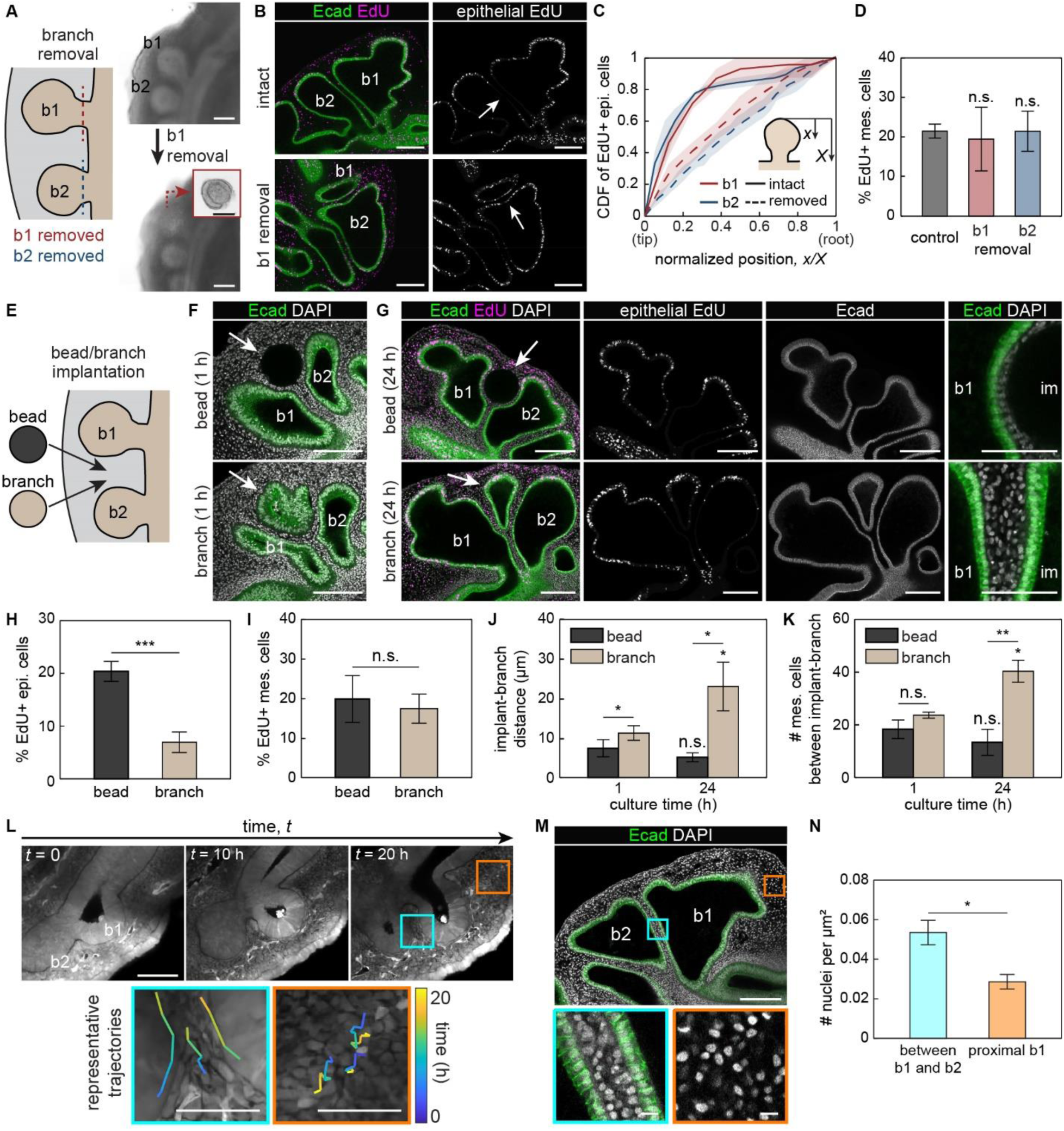
Surgically altering the spacing between branches affects the spatial patterning of epithelial proliferation. A. Schematic of experiment in which either b1 or b2 is surgically removed from the lung explant, and brightfield images of lung explant before and after removal of b1. Inset in the bottom image shows the surgically resected branch. Scale bars, 500 µm or 100 µm for insets. B. Fluorescence images of EdU incorporation in the epithelium of lung explants after removing b1, counterstained for E-cadherin (green). White arrows denote proximal stalk of b2. Scale bars, 100 µm. C. Graph showing the cumulative distribution function (CDF) of EdU-positive epithelial cells in b2 (b1 removal) and b1 (b2 removal), with or without surgical removal of the neighboring branch. Shown are the average and s.d. of 3 lungs per condition. D. Graph showing the percentage of EdU-positive cells in the mesenchyme. n.s. stands for not significant. E. Schematic of experiment in which either a bead or branch is surgically implanted into a recipient lung explant. F. Fluorescence images of nuclei (gray) in lung explants counterstained for E-cadherin (green). Shown are images of lungs after 1-hour culture after bead (top) or branch (bottom) implantation. White arrows denote the implanted bead/branch. Scale bars, 100 µm. G. Fluorescence images of EdU incorporation in the epithelium and E-cadherin staining of lung explants after bead (top) or branch (bottom) implantation. *E*6 lung explants were cultured for 24 hours after implantation. White arrows denote the implanted bead/branch. Rightmost images show the distances between implanted bead or branch and native branch (b1). im in the images denote either implanted bead or branch. Scale bars, 100 µm or 50 µm (inset). H. Graph showing the percentage of EdU-positive cells in the epithelium. Shown are the average and s.d. of 3 lungs per condition. ***, *p*<0.001. I. Graph showing the percentage of EdU-positive cells in the mesenchyme. Shown are the average and s.d. of 3 lungs per condition. J. Graph showing the distance between implanted bead or branch and neighboring native branches after culture. Shown are the average and s.d. of 4 lungs per condition. *, *p*<0.05. K. Graph showing the number of mesenchymal cells within a 50-μm-thick region of interest in the spacing between implanted bead or branch and native branch. Shown are the average and s.d. of 4 lungs per condition. *, p<0.05. **, p<0.005. L. Time-lapse images of a lung explant forming b2. Orange and cyan boxes denote the regions proximal to b1 and between b1 and b2, respectively. Representative trajectories of mesenchymal cells are shown in insets. Scale bars, 100 µm or 50 µm for insets. M. Fluorescence images of nuclei (gray) in lung explants counterstained for E-cadherin (green). Orange and cyan boxes denote the regions proximal to b1 and between b1 and b2, respectively. Scale bars, 100 µm or 10 µm for insets. N. Graph showing area-normalized nuclear density in the mesenchyme. Shown are the average and s.d. of 3 lungs per timepoint. *, *p*<0.05.

To confirm these qualitative observations, we conducted quantitative image analysis. Specifically, we measured the density of EdU-positive epithelial cells and plotted the cumulative distribution function (CDF) along the length of the branch. In the contralateral control bronchi, the CDF shows a steep slope near the tip that plateaus in the stalk region, indicating that most of the proliferation is localized to the epithelial cells within the tip of the branch (**Fig. 2C**). In the surgically resected side, however, the CDF has a nearly uniform slope, indicating that EdU-positive cells are evenly distributed along the length of the branch. In contrast, surgically removing either branch has no effect on EdU incorporation in the nearby mesenchymal cells (**Fig. 2D**), which continued to proliferate at a relatively high rate.

Curiously, we found that the increase in epithelial proliferation adjacent to the surgical site coincided with the emergence of what appeared to be ectopic branches (**Fig. S2A, B**). The embryonic chicken lung epithelium initiates branches via apical constriction downstream of actomyosin contraction.^20^ Staining for F-actin and phosphorylated myosin light chain (pMLC) revealed an absence of actomyosin contraction at the apical side of the emerging ectopic branches, suggesting that these branches do not form via apical constriction (**Fig. S2C**). Instead, these results suggest that the increase in epithelial proliferation after surgery may induce ectopic branches via buckling morphogenesis, similar to what is observed in mesenchyme-free cultured murine lung epithelium^21^ and avian lungs implanted with a focal source of growth factor.^22^

### Surgically decreasing spacing between branches leads to recovery of original branch spacing

Our branch resection experiments show that increasing the distance between branches increases the proliferation rate of the epithelium without appreciably affecting that of the mesenchyme. To determine the effects of decreasing the distance between branches, we surgically isolated a branch from a donor lung and inserted it in between b1 and b2 in a recipient lung (**Fig. 2E, F**). As a control, we implanted a PBS-containing agarose bead. We then cultured these explants for 24 hours before labeling for EdU incorporation (**Fig. 2G**). We found that inserting a bead increases EdU incorporation in the adjacent epithelium (**Fig. 2H**) but not the mesenchyme (**Fig. 2I**), consistent with our branch resection experiments. In contrast, inserting a branch has no effect on proliferation in either tissue layer (compare **Fig. 2H, I** to **Fig. 1H, I**). Nonetheless, we found that the spacing between the donor and recipient branches increased over time and reached a separation distance comparable to that between native branches (compare **Fig. 2J** to **Fig. 1F**). This separation distance correlated with an increase in the number of mesenchymal cells between the donor and recipient branches (**Fig. 2K**). These data thus suggest that branches in close proximity to each other lead to a decrease in epithelial proliferation and a simultaneous increase in mesenchymal cell number between branches.

Given that mesenchymal number increased without a corresponding increase in mesenchymal proliferation, we hypothesized that mesenchymal cells might migrate into the gap between epithelial branches. To test this hypothesis, we conducted single-cell resolution timelapse imaging of lungs from cytoplasmic-GFP-expressing chicken embryos (**Fig. 2L, Supplementary Video S1**). Tracking mesenchymal cells revealed that those located far from the branch exhibited stochastic motion. In contrast, those located adjacent to the epithelium exhibited directed migration into the region between b1 and b2. These patterns of migration correlated with an increase in the density of mesenchymal cells between the branches (**Fig. 2M, N**). Branch spacing thus correlates with decreases in epithelial proliferation and simultaneous increases in mesenchymal density.

### Exogenous FGF10 signaling does not affect patterns of epithelial proliferation or branch spacing

There are at least two possible mechanisms that could result in the observed patterns of epithelial proliferation and mesenchymal migration in tissues adjacent to neighboring branches (**Fig. 3A**). First, since the mesenchyme is the main source of mitogenic growth factors in the developing lung,^5,23^ the small volume of this tissue between the two bronchi may limit the concentration of growth factors and thus reduce epithelial proliferation. Alternatively, since the epithelia of many branched organs produce inhibitory morphogens, the narrow spacing between the two bronchi may result in a locally high concentration of such inhibitors and reduce epithelial proliferation. Importantly, many of the molecules that act as mitogens or inhibitory morphogens in the epithelium have also been found to alter mesenchymal migration and condensation.^24–30^

**Fig. 3.**
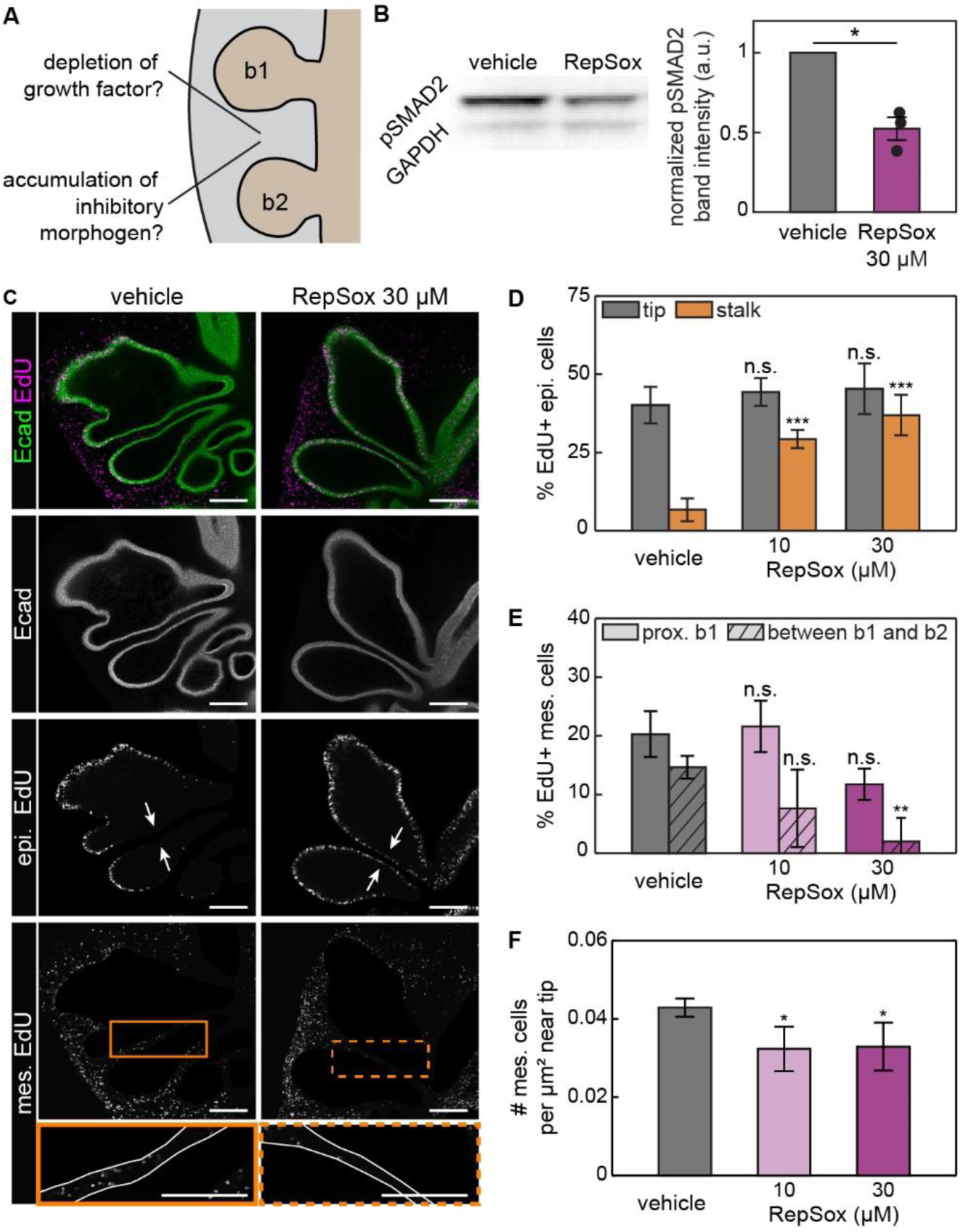
Inhibiting signaling from TGFβ disrupts the spatial pattern of proliferation and mesenchymal density in the developing lung. A. Schematic showing two possible mechanisms to spatially pattern proliferation in the embryonic lung. B. Representative immunoblots for pSMAD2 and GAPDH in lungs cultured with RepSox or vehicle control, and graph showing the normalized pSMAD2 band intensity from 3 replicates. Shown are the average and s.d. of normalized intensity of the pSMAD2 band. C. Fluorescence images of EdU incorporation in the epithelium and mesenchyme of lung explants cultured with RepSox or vehicle control. White arrows denote closely apposed branch stalks. Solid and dotted boxes denote the regions between b1 and b2, respectively. Scale bars, 100 µm. D. Graph showing the percentage of EdU-positive epithelial cells in lung explants cultured with RepSox or vehicle control. Shown are the average and s.d. of 6 lungs per condition. n.s. stands for not significant. \*\*\**p*<0.001. E. Graph showing the percentage of EdU-positive mesenchymal cells in lung explants cultured with RepSox or vehicle control. Shown are the average and s.d. of 3 lungs per condition. \*\**p*<0.005. F. Graph showing the density of mesenchymal cells near the tip in lung explants cultured with RepSox or vehicle control. Shown are the average and s.d. of at least 4 lungs per condition. \**p*<0.05.

We thus conducted spatial transcriptomics^13–15^ on *E*5 lungs to identify gene-expression patterns that are associated with each possible mechanism. Our transcriptomics data show that the mesothelium expresses *Fgf10*, which encodes for fibroblast growth factor 10 (FGF10), a key regulator of epithelial proliferation in the developing lung (**Fig. S3A**).^23,30,31^ In contrast, the gene that encodes for its receptor, *Fgfr2*, is highly expressed in both the epithelium and subepithelial mesenchyme (**Fig. S3A**). These spatial patterns of expression were confirmed by fluorescence in situ hybridization (RNAscope) analysis (**Fig. S3B**). We then tested whether the reduction in proliferation in the epithelium adjacent to a neighboring branch results from depletion of FGF10. Specifically, we supplemented the culture media with exogenous FGF10 to increase the concentration surrounding epithelial branches. We first confirmed that exogenous FGF10 activates the FGF signaling pathway by immunoblotting for phosphorylated ERK (pERK)1/2^32^ in lungs cultured with or without FGF10 (**Fig. S3C**). Despite the increase in pERK1/2, we found that adding exogenous FGF10 has no effect on the pattern of epithelial proliferation: we still observed a decrease in the percentage of EdU-positive cells in the stalks adjacent to neighboring branches (**Fig. S3D, E**). Furthermore, the ratio of EdU-positive mesenchymal cells (**Fig. S3F**) and density of mesenchymal cells (**Fig. S3G**) was similar to that in vehicle-treated controls. These data suggest that patterns of proliferation and mesenchymal migration within and around elongating branches are unlikely to result from depletion of FGF10.

### Blocking TGFβ signaling alters patterns of epithelial proliferation

We next tested whether the patterns of epithelial proliferation and mesenchymal migration result from signaling downstream of inhibitory morphogens (**Fig. 3A**). Disrupting signaling from BMP or TGFβ causes neighboring branches to contact each other in cultured kidney explants.^9^ Spatial transcriptomics (**Fig. S4A-D**) and fluorescence in situ hybridization analysis (**Fig. S4E, F**) revealed that both BMP and TGFβ ligands as well as their receptors are expressed at these stages of development in the embryonic chicken lung. We therefore tested whether the proliferation and spacing of epithelial branches are downstream of inhibitory signaling from BMP or TGFβ.

First, we disrupted BMP signaling by supplementing the culture media with dorsomorphin, a small molecule that inhibits the BMP type I receptor,^33^ as confirmed by immunofluorescence analysis for pSMAD1/5/9 (**Fig. S5A, B**).^34^ Lungs cultured in the presence of dorsomorphin showed similar patterns of proliferation as vehicle-treated controls, with a high percentage of EdU-positive epithelial cells at the tips of the branches and lower levels in stalks adjacent to neighboring branches (**Fig. S5A, C**). Proliferation (**Fig. S5D**) and density (**Fig. S5E**) in the mesenchyme were also similar in dorsomorphin-treated explants and vehicle-treated controls. These data suggest that BMP signaling is not required to specify patterns of proliferation or migration within and around elongating branches.

We then disrupted TGFβ signaling by supplementing the culture media with RepSox, a small molecule that inhibits the type I TGFβ receptor,^35,36^ as confirmed by immunoblotting for pSMAD2 (**Fig. 3B**). Explants of *E*6 lungs were cultured for 24 hours in the presence of RepSox, which led to a striking increase in the percentage of EdU-positive epithelial cells in the stalks adjacent to neighboring branches (**Fig. 3C, D**). Importantly, the percentage of EdU-positive epithelial cells at the tip of the branch was unaffected by the presence of the inhibitor (**Fig. 3C, D**). The mesenchyme was also affected by RepSox, which led to an overall decrease in the percentage of EdU-positive cells (**Fig. 3C, E**). This effect was especially stark in between b1 and b2, where almost no EdU-positive mesenchymal cells were observed in lungs treated with a high concentration of RepSox. Mesenchymal cell density in between the tips also decreased with RepSox treatment (**Fig. 3F**). The patterns of proliferation and mesenchymal density within and between branches in the embryonic chicken lung thus appear to depend on signaling downstream of TGFβ.

### Mesenchymal condensations form near sources of TGFβ and can displace the adjacent epithelium

Our data indicate that closely apposed epithelial branches are inhibited from proliferation downstream of TGFβ signaling. Such paracrine interactions are widely observed across branching organs and are thought to be the primary mechanism by which epithelia avoid each other. However, our data also suggest that these patterns of epithelial proliferation alone cannot explain the spacing between branches. First, although a decrease in proliferation within neighboring stalks is first detected at *E*7, the regular spacing between branches is already established at *E*6 (**Fig. 1F**). Second, surgically implanting a donor branch leads to recovery of branch spacing without affecting epithelial proliferation (**Fig. 2H-J**).

We therefore focused more closely on how TGFβ signaling affects mesenchymal dynamics during branch elongation. To define the effects of TGFβ, we implanted TGFβ-containing agarose beads in the mesenchyme between the tips of b1 and b2 in *E*6 lung explants and cultured them for 24 hours (**Fig. 4A**). EdU analysis confirmed that TGFβ locally inhibits epithelial proliferation adjacent to the bead (**Fig. S6A**).^37–39^ In lungs implanted with vehicle-containing beads, the epithelial branches extended and nearly contacted the bead (**Fig. 4B, C**). In lungs implanted with TGFβ-containing beads, however, the distance between the epithelium and the bead increased over time (**Fig. 4B, C**), similar to explants in which we implanted donor branches (**Fig. 2E, J**). Moreover, the epithelium also exhibited a concentric shape around the mesenchyme surrounding the bead, maintaining a nearly uniform distance at any given point (**Fig. 4B**), reminiscent of the parallel spacing between branches in the lung. Consistently, treatment with RepSox blocked changes in branch spacing when we implanted donor branches into recipient lungs (**Fig. S6B**). These observations confirm that, in addition to affecting patterns of proliferation, TGFβ affects the thickness of the mesenchymal layer between epithelial branches.

**Fig. 4.**
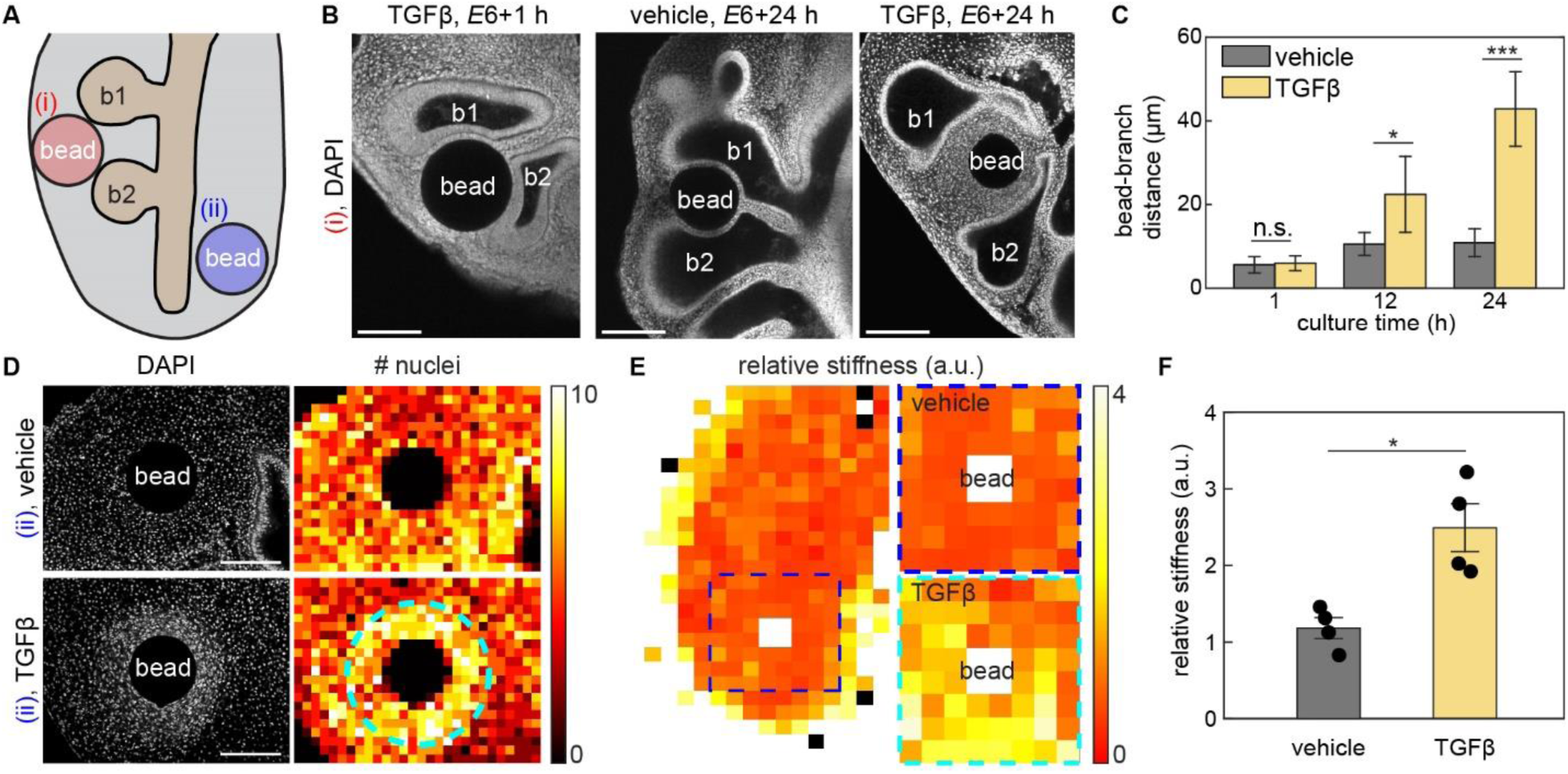
Implanting a focal source of TGFβ induces the formation of mesenchymal condensations, which physically displace the adjacent epithelium. A. Schematic of bead-insertion experiment in which a bead is inserted either (i) in between b1 and b2 or (ii) near the non-branching ventral epithelium. B. Fluorescence images of nuclei in lung explants in which beads were inserted in between b1 and b2. Shown are images of lungs after 1-hour (TGFβ, left) or 24-hour culture (vehicle, middle; TGFβ, right). Scale bars, 100 µm. C. Graph showing the bead-branch distance over time in lung explants implanted with TGFβ-containing beads or vehicle-containing controls. Shown are the average and s.d. of 3 lungs per condition. *, *p*<0.05. ***, *p*<0.001. D. Fluorescence images of nuclei in lung explants implanted with either a TGFβ-containing bead or vehicle-containing control. Heatmaps show nuclear density, and the dotted line in the heatmap of the lung implanted with a TGFβ-containing bead denotes the boundary of the mesenchymal condensation. Scale bars, 100 µm. E. Representative heatmap showing stiffness of the mesenchyme in a bead-implanted lung. The dotted line denotes the region adjacent to the implanted bead, and representative heatmaps near the TGFβ-containing bead or vehicle-containing control are shown. Measurements are normalized to the stiffness of the mesenchyme near ventral airway. F. Graph showing the stiffness of the mesenchyme near the implanted bead. Shown are the average and s.d., and each dot represents one embryo. *, *p*<0.05.

We next examined the effects of TGFβ on mesenchymal cells independent of epithelial branches. Specifically, we implanted TGFβ-containing beads adjacent to the non-branching ventral airway epithelium (**Fig. 4A**).^22^ Nuclear staining revealed an increase in mesenchymal cell density in a concentric pattern around TGFβ-containing beads as compared to vehicle-containing controls (**Fig. 4D**). Increasing the concentration of TGFβ loaded into the beads increased the size of the mesenchymal condensation (**Fig. S6C, D**), which exhibited strong pSMAD2 staining (**Fig. S6E**) and coincided with an increase in the local density of perlecan (**Fig. S6F**), the deposition of which is known to increase in response to TGFβ.^40,41^ Furthermore, the mesenchyme adjacent to TGFβ-containing beads was stiffer than that adjacent to vehicle-containing beads (**Fig. 4E, F**), consistent with formation of a condensation in the former. Adding RepSox to the culture media was sufficient to prevent mesenchymal condensation around TGFβ-containing beads (**Fig. S6G**). The spherical shape of the mesenchymal condensations matched the concentric shape of the epithelium around TGFβ-containing beads (**Fig. 4B**), suggesting that the airway epithelium is shaped by physical interactions with the adjacent mesenchyme, which changes density and stiffness in response to TGFβ.

### TGFβ-induced condensation of mesenchymal cells is driven by directed migration

We therefore investigated the mechanism by which the mesenchyme condensed. Mesenchymal condensations can result from a local increase in proliferation and/or from directed cell migration.^42^ TGFβ is a known activating morphogen for mesenchymal cells,^43,44^ and strong pSMAD2 staining is observed in subepithelial mesenchymal cells (**Fig. S6H**). Consistently, our experimental manipulations showed that the mesenchyme is both proliferative and migratory at this stage of development. To parse the relative effects of proliferation and directed migration in TGFβ-induced condensation of mesenchymal cells and epithelial displacement, we used agent-based modeling.^16,17^

In our computational model, each cell is treated as a single agent that interacts with its neighbors (**Fig. 5A**). Specifically, mesenchymal cells exhibit Brownian motion by default, but their movements are affected by interaction with adjacent cells, which acts as a repulsive force when the distance between two cells becomes too small. In contrast, epithelial cells are connected to each other to model the branch, and their behaviors are governed by epithelial tension, luminal pressure, and epithelial bending in addition to interaction forces.

**Fig. 5.**
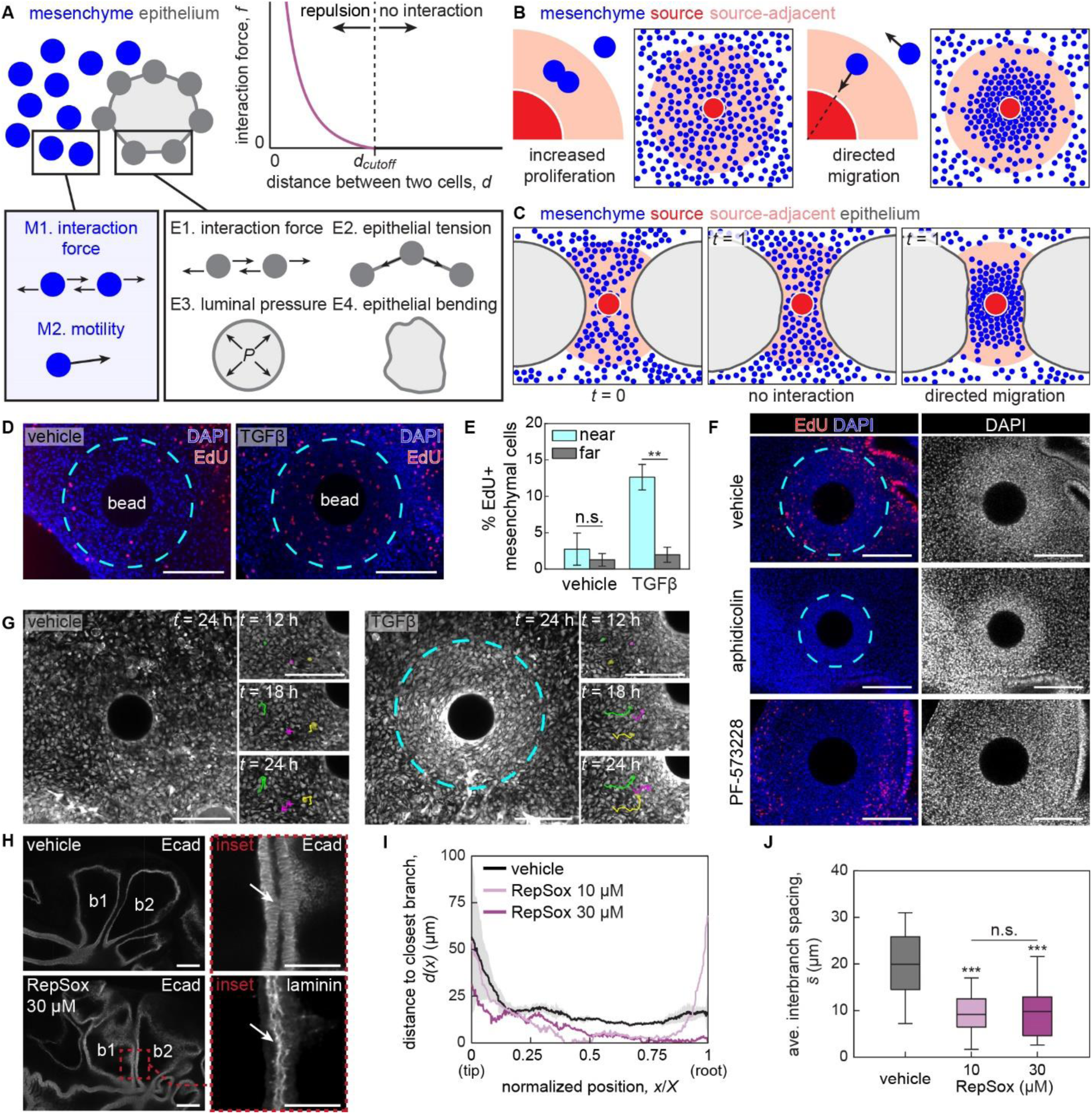
Directed migration results in TGFβ-induced mesenchymal condensation. A. Schematic of the parameters used to model the epithelium and mesenchyme. B. Simulation results showing that directed migration, but not increased proliferation, leads to the formation of mesenchymal condensations. C. Simulation results showing that mechanical interactions between the mesenchymal condensation and adjacent epithelium can lead to branch deformation. D. Fluorescence images of EdU incorporation (red) in lung explants implanted with TGFβ-containing bead or vehicle-containing control, counterstained for nuclei (blue). Cyan dotted lines denote the boundary of the condensation (right) and an identical-sized circle in the vehicle control (left). Scale bars, 100 µm. E. Graph showing the percentage of EdU-positive cells in the mesenchyme near to and far from the bead in lungs implanted with TGFβ-containing beads or vehicle-containing controls. Blue bars denote the ratio in cells located within the condensation (TGFβ) or within a region of equivalent average size (vehicle). Shown are the average and s.d. of 3 lungs per condition. F. Fluorescence images of EdU incorporation (red) in lung explants inserted with TGFβ-containing beads and treated with vehicle control, aphidicolin, or PF-573228, counterstained for nuclei (blue). Scale bars, 100 µm. G. Live-imaging of GFP-expressing lungs implanted with either a TGFβ-containing bead or vehicle-containing control. Time-lapse images showing representative trajectories of mesenchymal cells near the bottom-left quadrant of the bead are shown together. Scale bars, 100 µm. H. Fluorescence images of immunofluorescence staining for laminin (magenta) and E-cadherin (green) in lung explants cultured with RepSox or vehicle control. White arrows in the inset show site of contact between b1 and b2. Scale bars, 100 µm. I. Graph showing the distance between b1 and b2 in lungs cultured with RepSox or vehicle control. Vehicle control is the average of 3 samples, while representative data are shown for experimental conditions. J. Graph of average spacing between branches in lungs cultured with RepSox or vehicle control. Shown are average and s.d. of 3 lungs per condition. n.s., not significant. ***, *p*<0.001.

We first tested the role of proliferation and directed migration in mesenchymal condensation. We introduced a circular source of activating morphogen, which represents a TGFβ-containing bead, and either assumed enhanced proliferation or directed migration of mesenchymal cells toward the source (**Fig. 5B**). While enhanced proliferation did not result in an appreciable change in the density of mesenchymal cells, directed migration resulted in a condensation-like, densely packed mesenchyme surrounding the bead (**Fig. 5B**). Therefore, our computational model suggests that directed migration, but not increased proliferation, is required for mesenchymal cells to condense around a source of TGFβ.

We then used our computational model to place the morphogen source in between two epithelial branches and tested whether the physical interactions between a mesenchymal condensation and its adjacent epithelium can lead to branch deformation (**Fig. 5C**). In the absence of directed migration, which represents mesenchymal response to a vehicle-containing control bead (**Fig. 4B**), epithelial branches continue to expand and decrease the distance between them and the bead over time. In contrast, in the presence of directed migration, a mesenchymal condensation forms near the bead and the adjacent epithelium deforms around the condensation. The resulting epithelial morphology is comparable to what we observed experimentally in TGFβ-containing bead-implanted lung explants (**Fig. 4B**), suggesting that physical interactions between the mesenchyme and epithelium can affect the morphology of the latter.

We then experimentally tested the predictions obtained from our agent-based model. EdU analysis revealed enhanced proliferation within mesenchymal condensations (**Fig. 5D, E**). However, inhibiting proliferation using aphidicolin had no effect on the formation of condensations (**Fig. 5F**). Single-cell resolution live-imaging of lungs explanted from cytoplasmic-GFP-expressing chicken embryos revealed convergent motion of mesenchymal cells toward TGFβ-containing beads, but not toward vehicle-containing controls (**Fig. 5G, Supplementary Video S2, S3**).^29^ Consistently, inhibiting cell migration using the focal adhesion kinase inhibitor, PF-573228, prevented the formation of mesenchymal condensations (**Fig. 5F**). These data reveal that TGFβ promotes mesenchymal proliferation and directed migration, but only the latter is necessary for the formation of condensations, confirming the predictions of our model.

### Continuous inhibition of TGFβ signaling leads to branch-branch contact

If densely packed mesenchymal cells maintain the spacing between epithelial branches, then we would expect that preventing mesenchymal condensation might permit epithelial branches to widen and eventually contact their neighbors. We therefore tested whether continuously disrupting TGFβ signaling would promote lateral growth and result in contact between branches. To that end, we cultured *E*6 lung explants in RepSox-containing culture media for 72 hours and compared the shape of the epithelium with that of vehicle-treated controls. As expected, we found that the stalks of neighboring branches were aligned parallel to each other in control explants (**Fig. 5H, I**), separated by an average distance of ∼20 μm (**Fig. 5J**). In contrast, in RepSox-treated explants, the stalks of neighboring branches were not parallel, showed a reduction in spacing (**Fig. 5J**), and frequently contacted each other (**Fig. 5H, I**).

Fluorescently labeling nuclei confirmed that the contacting epithelia lacked mesenchymal cells between them. Surprisingly, however, immunofluorescence for pan-laminin revealed that the contacting branches were still separated by a basement membrane (**Fig. 5H**). These data suggest that TGFβ signaling regulates both epithelial and mesenchymal cell behaviors to control the spatial patterns of proliferation, cell migration, and spacing between branches in the embryonic chicken lung.

### TGFβ induces mesenchymal condensation in embryonic mouse lungs and salivary glands, but not in kidneys

We then asked whether the cellular responses to TGFβ in the embryonic chicken lung are conserved in other branched organs. To answer that question, we implanted TGFβ-containing beads into the mesenchyme of kidney, lung, and salivary gland explants from mouse embryos, cultured them for 24 hours (**Fig. 6A**), and then measured the distance between each bead and its closest epithelial branch.

**Fig. 6.**
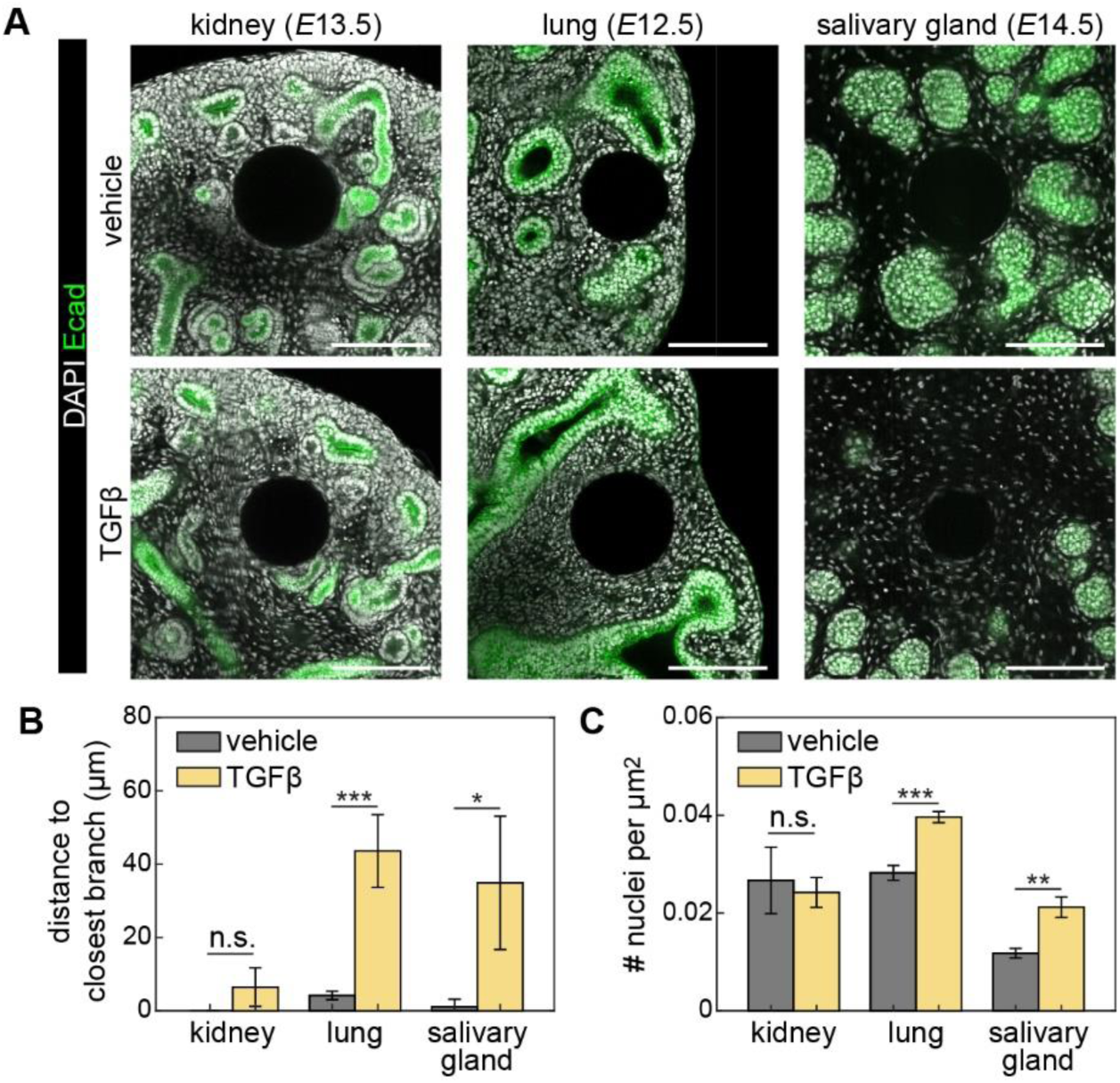
TGFβ-induced mesenchymal condensation in branched organs from mouse embryos. A. Fluorescence images of nuclei (gray) in kidney, lung, and salivary gland explants counterstained for E-cadherin (green). Scale bars, 100 µm. B. Graph showing distance to closest branch from the implanted bead. Shown are the average and s.d. of at least 3 explants per timepoint. n.s., not significant. *, *p*<0.05, ***, *p*<0.001. C. Graph showing area-normalized nuclear density in the mesenchyme near the bead. Shown are the average and s.d. of 3 explants per timepoint. **, *p*<0.005.

In the kidney, implanting TGFβ-containing beads produced results similar to that of vehicle-containing controls. E-cadherin-stained epithelium frequently contacted both TGFβ- and vehicle-containing beads, and the measured mesenchymal cell density was comparable in both conditions (**Fig. 6B, C**). These observations are consistent with the fact that the developing kidney uses BMP, rather than TGFβ, to achieve epithelial self-avoidance.^9^ In the developing lung and salivary gland, however, implantation of TGFβ-containing beads led to an increase in both the distance to the closest epithelial branch and the density of adjacent mesenchymal cells (**Fig. 6B, C**). These observations indicate that TGFβ-induced mesenchymal cell condensation might operate across organs to tune the spacing between epithelial branches.

## Discussion

Taking advantage of the uniform spacing within the embryonic chicken lung allowed us to elucidate how epithelial branches avoid each other during embryonic development. Epithelial cells in close proximity to neighboring branches showed a reduction in proliferation, a pattern that was lost when the neighboring branch was surgically removed. Supplementing the culture media with FGF10 or blocking BMP signaling had no effect on the pattern of proliferation. However, blocking TGFβ signaling significantly increased the percentage of proliferating epithelial cells. Mesenchymal cells also responded to TGFβ by forming a stiff, local condensation in a concentration-dependent manner by directed migration, which affected the spacing between branches by physically displacing the nearby epithelium. Continuous inhibition of TGFβ prevented mesenchymal condensation and eventually resulted in branches contacting each other. These data suggest that the spacing of the branches within the embryonic chicken lung depends on physical interactions between the epithelium and mesenchyme in response to this morphogen.

TGFβ has been shown to act as an inhibitory morphogen in other branched epithelia, including the mammary and salivary glands. In the mammary gland, delivering exogenous TGFβ by implanting TGFβ-containing pellets or beads resulted in regression of end buds and inhibition of ductal growth, effects that were reversed upon removal of the implant.^8,18^ Moreover, overexpression of TGFβ1 in mouse mammary epithelial cells inhibited branching, while disrupting TGFβ signaling resulted in increased branching.^45^ In the salivary gland, adding exogenous TGFβ1 inhibited epithelial growth and branching in cultured explants.^46^ However, inhibiting signaling downstream of the type I TGFβ receptor was not sufficient to disrupt self-avoidance of the salivary gland epithelium.^10^ Nonetheless, here we found that implanting TGFβ-containing beads led to the formation of mesenchymal condensations and altered epithelial shape in the mouse embryonic salivary gland and lung, but not the kidney. These data suggest that mesenchymal condensations may be conserved in a subset of branching organs. It is worth noting that while previous studies have primarily focused on the inhibitory effects of TGFβ on epithelial cells, our data suggest that its activating effects on mesenchymal cells are equally, if not more, important for achieving branch spacing.

Explant cultures of several branched organs have suggested that epithelial branches are capable of “sensing” neighboring branches when the distance between them reaches ∼30 μm, as the spacing does not decrease further.^9,10,18,19,47,48^ Here, we found that the size of the mesenchymal condensations that form in response to TGFβ control the spacing between branches by acting as a physical barrier in the developing chicken lung. TGFβ was previously found to promote mesenchymal condensation and chondrogenesis in chicken embryos.^43,44,49^ However, how TGFβ affects the cellular dynamics of lung mesenchymal cells, and how mesenchymal condensations physically interact with adjacent epithelial branches, has not been previously reported to the best of our knowledge. We further found that condensation size is a function of TGFβ concentration, and that spacing between innate and implanted branches recovers that of intact lungs within 24 hours of culture. Together, these results suggest that the regular spacing observed during development may arise from the coordinated responses of epithelial and mesenchymal cells to the local concentration of morphogens such as TGFβ, which act differentially on the two tissue types to determine cell density and branch spacing.

Directed migration of mesenchymal cells is crucial for the formation of condensations during tooth development and chondrogenesis.^27,50^ However, different organs use different molecular signals to induce such directed migration. In the developing tooth, the dental epithelium produces FGF8, which acts as an activating morphogen that attracts mesenchymal cells.^27^ In the developing cartilage, local deposition of fibronectin at the center of condensations, which is sometimes induced by TGFβ, recruits mesenchymal cells.^50^ We observed upregulation of perlecan deposition near implanted TGFβ-containing beads, which may suggest a similar role for the ECM in promoting, refining, or maintaining mesenchymal condensations in the developing lung.

Our findings demonstrate that diffusion of morphogens regulates the spacing between branches and prevents them from colliding with each other in the developing chicken lung by affecting both the epithelium and mesenchyme. Such regulated spacing is ubiquitous across branched organs and critical for maximizing the area of the material-exchanging interface.

Whether these organs exploit identical molecular mechanisms to regulate the spacing between branches is an interesting question that remains to be answered. In particular, if the size of mesenchymal condensations determines branch spacing, it will be interesting to determine whether a similar mechanism can also work in organs where the epithelium is surrounded by relatively larger and more densely packed cells, such as the mammary gland. On the other hand, the epithelial branches of the bird lung merge with each other at later stages of development to form a continuous circuit for airflow.^51^ How the mesenchyme surrounding epithelial branches in the embryonic lung shift its behavior from blocking to permitting epithelial contact also requires further investigation.

## Abbreviations

BMP: bone morphogenetic protein;
FGF: fibroblast growth factor;
HH: Hamburger-Hamilton;
pMLC: phosphorylated myosin light chain;
TGF: transforming growth factor;

## Acknowledgements

We thank the Tissue Morphodynamics Group for helpful discussions, and G. Laevsky and S. Wang for assistance with imaging. This work was supported in part by grants from the National Institutes of Health (NIH; HD099030, HD111539, HL164861, HL166311) and the National Science Foundation (2134935). C.J.P. was supported in part by a postdoctoral fellowship from the American Heart Association (25POST1375169). P.Z. was supported in part by a Princeton Bioengineering Initiative - Innovators (PBI2), Distinguished Postdoctoral Fellowship. CT-Y was supported in part by the New Jersey Department of Health, the Division of Office of Research Initiatives, and the New Jersey Commission on Cancer Research (NJCCR) through a 2023 NJCCR Postdoctoral Research Grant, as well as the Damon Runyon Cancer Research Foundation through the Quantitative Biology Postdoctoral Fellowship (DRQ-17-23).

## Supplementary Figure Legends

**Fig. S1.**
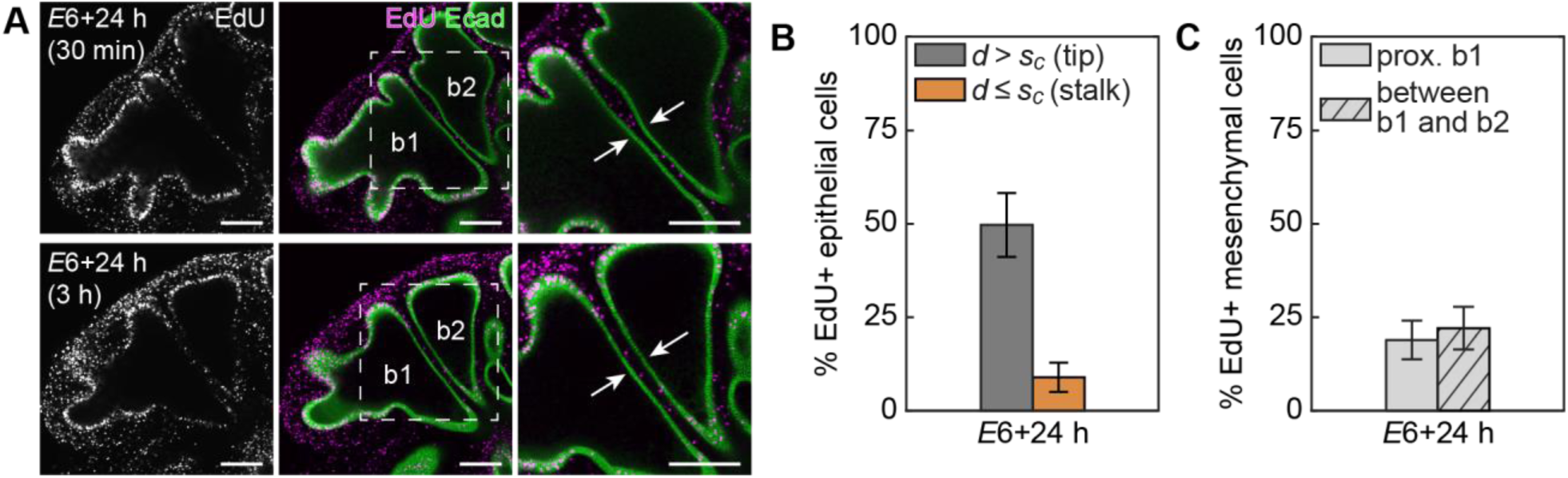
EdU analysis in the epithelium and mesenchyme of cultured lung explants – related to Fig. 1. (**A**) Fluorescence images of EdU incorporation (magenta) in embryonic chicken lung explants counterstained for E-cadherin (green), with either 30 min or 3 hours of EdU pulse. Scale bars, 100 μm. Graph of percentage of EdU-positive cells in the (**B**) epithelium and (**C**) mesenchyme of cultured lung explants. In panel (**C**), solid and hatched bars denote percentage of EdU-positive mesenchymal cells proximal to b1 and in between b1 and b2, respectively.

**Fig. S2.**
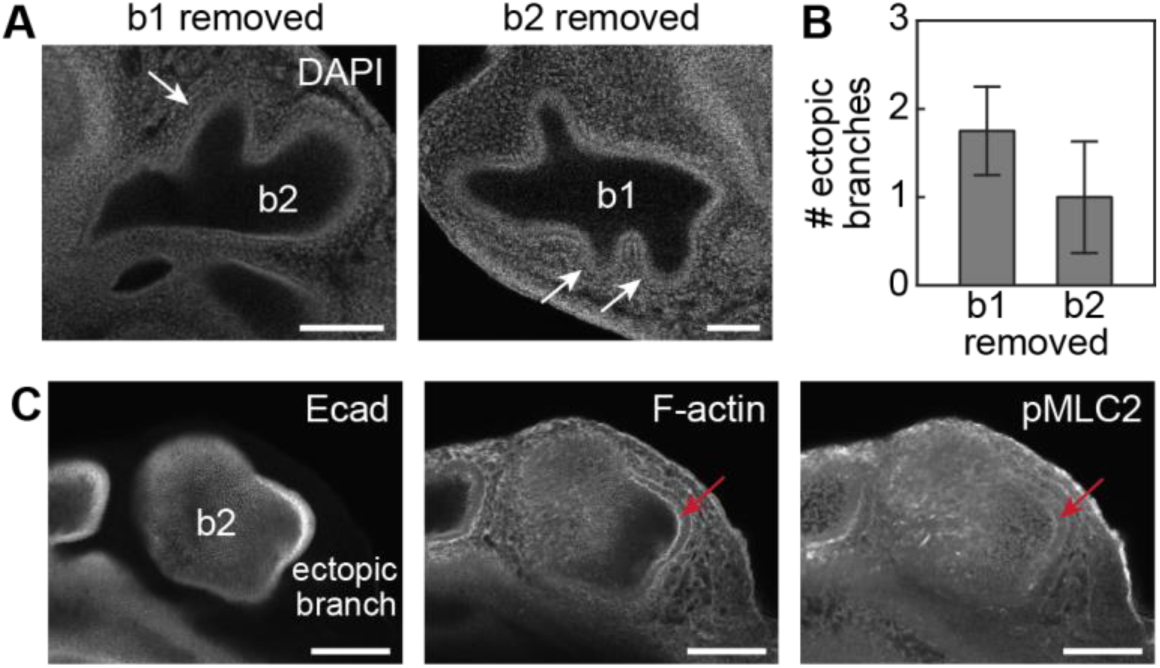
Analysis of apical constriction in ectopic branches – related to Fig. 2. (**A**) Fluorescence images of nuclei in lung explants after surgical removal of b1 or b2, focused on the remaining neighboring branch. Scale bars, 100 μm. (**B**) Graph showing number of ectopic branches that form in the remaining neighboring branch after surgical removal of b1 (i) or b2 (ii). Shown are average and s.d. of at least 4 lungs per condition. (**C**) Confocal section of an ectopic branch stained for F-actin and pMLC2 lacks noticeable signal on the apical side. Scale bars, 100 μm.

**Fig. S3.**
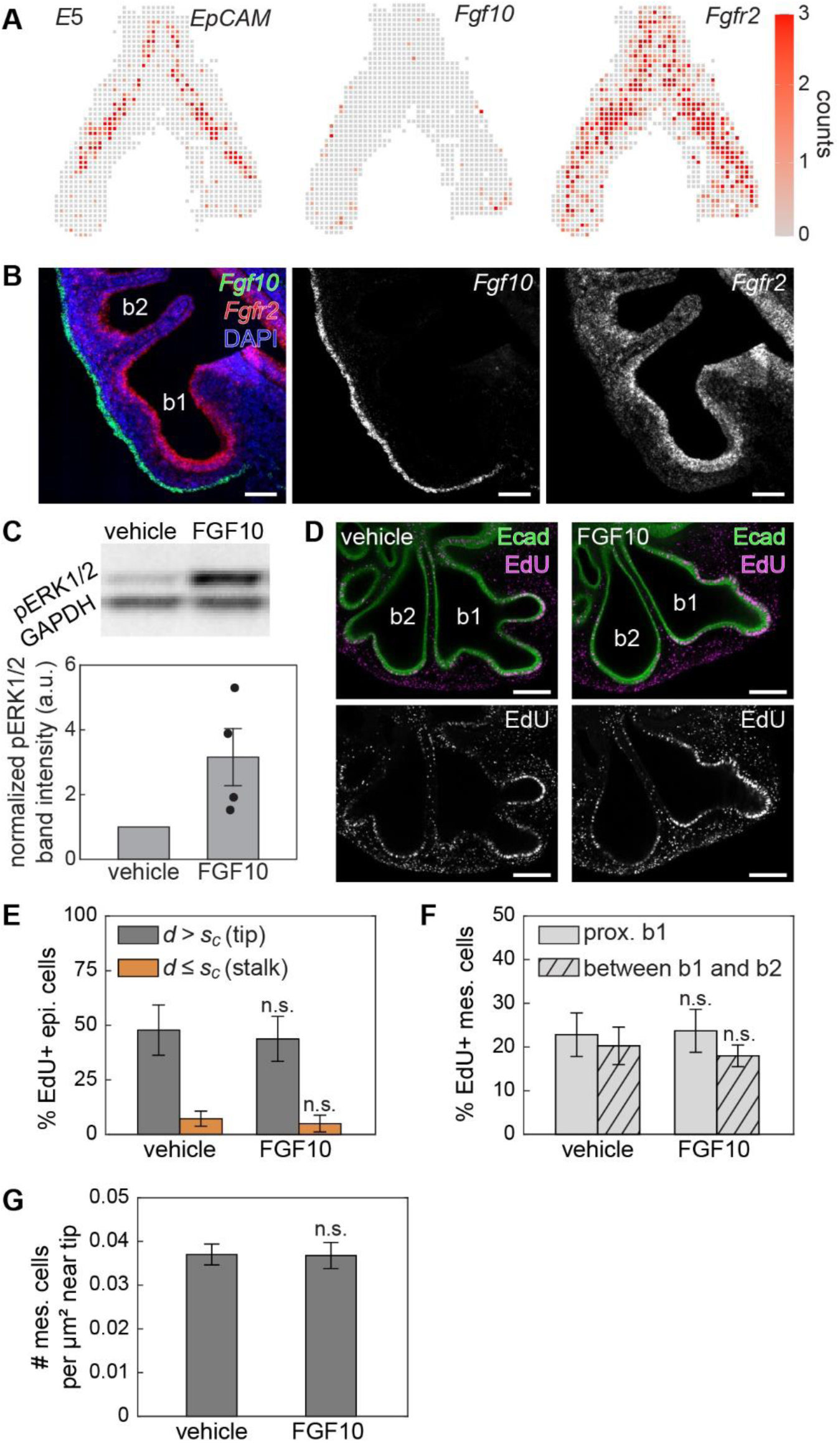
Exogeneous addition of FGF10 has no effect on the spatial pattern of proliferation in the embryonic lung – related to Fig. 3. (**A**) Representative gene-expression patterns of *EpCAM*, which marks the epithelium, *Fgf10,* and *Fgfr2*. (**B**) In situ hybridization analysis for *Fgf10* and *Fgfr2* in *E*6 lungs cultured for 24 hours. Scale bars, 100 µm. (**C**) Representative immunoblots for pERK1/2 and GAPDH in lungs cultured with FGF10 or vehicle control, and graph showing the normalized intensity of the pERK1/2 band from 4 replicates. (**D**) Fluorescence images of EdU incorporation (magenta) in lung explants cultured with exogeneous FGF10 or vehicle control, counterstained for E-cadherin (green). Scale bars, 100 µm. (**E**) Graph showing the percentage of EdU-positive cells in the epithelium with and without exogenous FGF10. Shown are the average and s.d. of 3 lungs per condition. (**F**) Graph of percentage of EdU-positive cells in the mesenchyme in lungs cultured with FGF10 or vehicle control. Solid and hatched bars denote percentage of EdU-positive mesenchymal cells proximal to b1 and in between b1 and b2, respectively. n.s. stands for not significant. (**G**) Graph showing the density of mesenchymal cells near the tip in lung explants cultured with exogenous FGF10 or vehicle control. Shown are the average and s.d. of 3 lungs per condition.

**Fig. S4.**
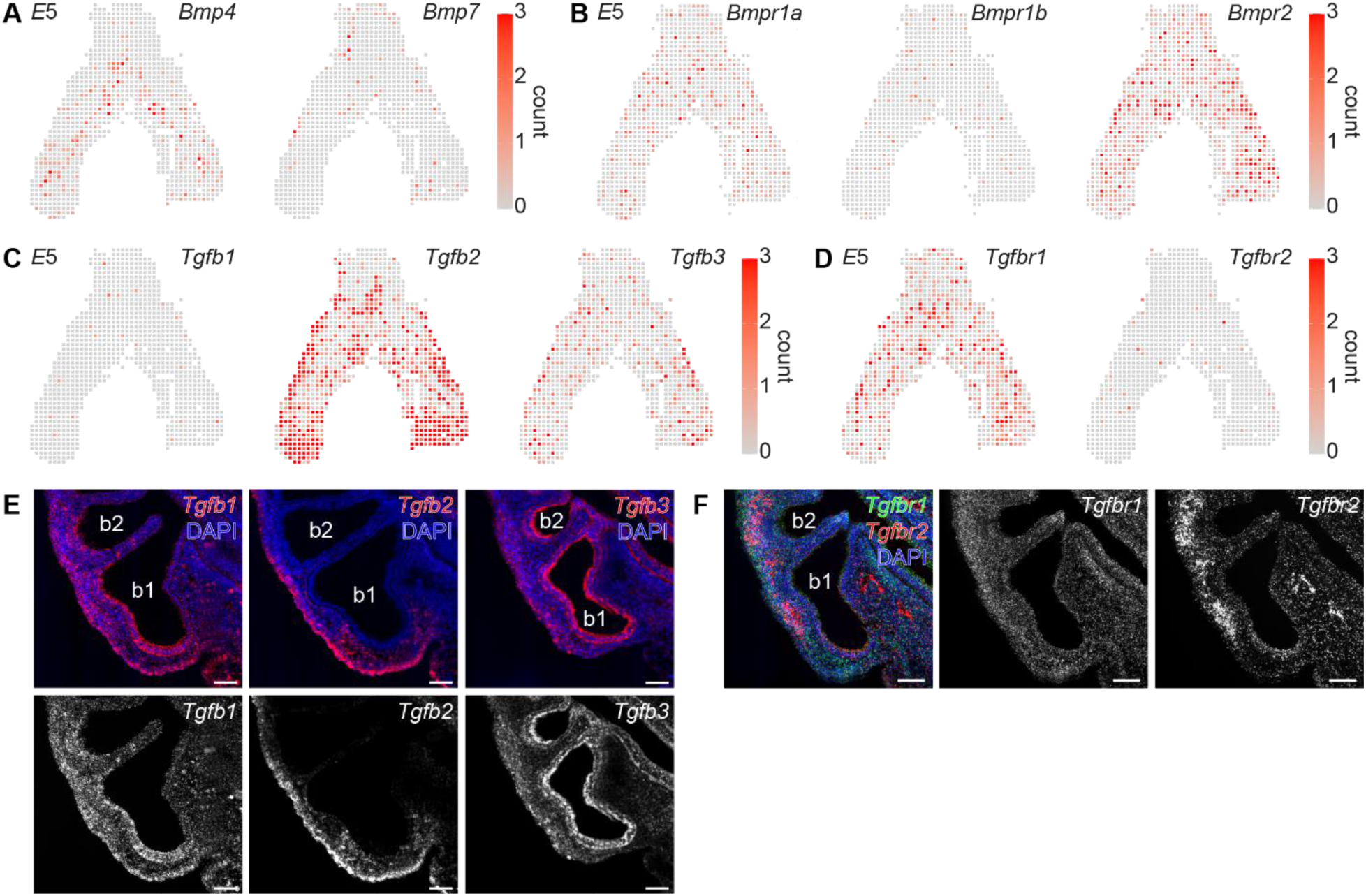
Spatial patterns of expression of inhibitory morphogens for epithelial cells and their receptors in the developing lung – related to Fig. 3. (**A**) Representative gene-expression patterns of *Bmp4* and *Bmp7*. (**B**) Representative gene-expression patterns of *Bmpr1a*, *Bmpr1b*, and *Bmpr2*. (**C**) Representative gene-expression patterns of *Tgfb1*, *Tgfb2*, and *Tgfb3*. (**D**) Representative gene-expression patterns of *Tgfbr1* and *Tgfbr2*. (**E**) Fluorescence in situ hybridization for *Tgfb1*, *Tgfb2*, and *Tgfb3* in *E*6 lungs cultured for 24 hours. Scale bars, 100 µm. (**F**) Fluorescence in situ hybridization for *Tgfbr1* and *Tgfbr2* in *E*6 lungs cultured for 24 hours. Scale bars, 100 µm.

**Fig. S5.**
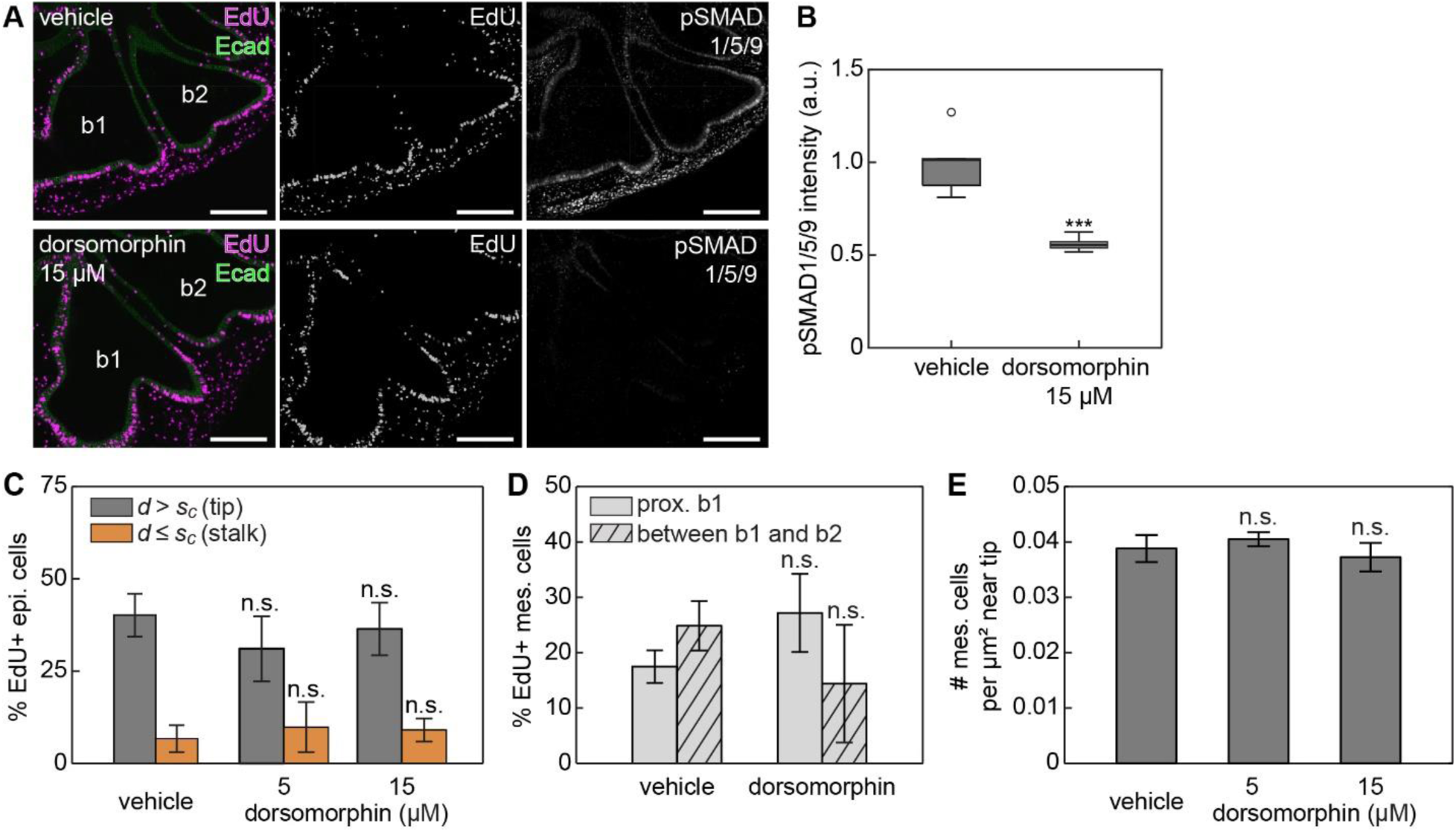
Inhibiting signaling from BMP has no effect on the spatial patterns of proliferation in the embryonic lung – related to Fig. 3. (**A**) Fluorescence images of EdU incorporation (magenta) and pSMAD1/5/9 staining in lung explants cultured with BMP receptor inhibitor, dorsomorphin, or vehicle control, counterstained for E-cadherin (green). Scale bars, 100 µm. (**B**) Graph showing the relative intensity of pSMAD1/5/9 staining in lung explants cultured with dorsomorphin or vehicle control. Shown are the average and s.d. of 3 lungs per condition. \*\*\**p*<0.001 (**C**) Graph showing the percentage of EdU-positive cells in the epithelium of lung explants cultured with dorsomorphin or vehicle control. Shown are the average and s.d. of 6 lungs per condition. n.s. stands for not significant. (**D**) Graph showing the percentage of EdU-positive cells in the mesenchyme in lungs cultured with dorsomorphin or vehicle control. Solid and hatched bars respectively denote percentage of EdU-positive mesenchymal cells proximal to b1 and in between b1 and b2. n.s. stands for not significant. (**E**) Graph showing the density of mesenchymal cells near the tip in lung explants cultured with dorsomorphin or vehicle control. Shown are the average and s.d. of 3 lungs per condition.

**Fig. S6.**
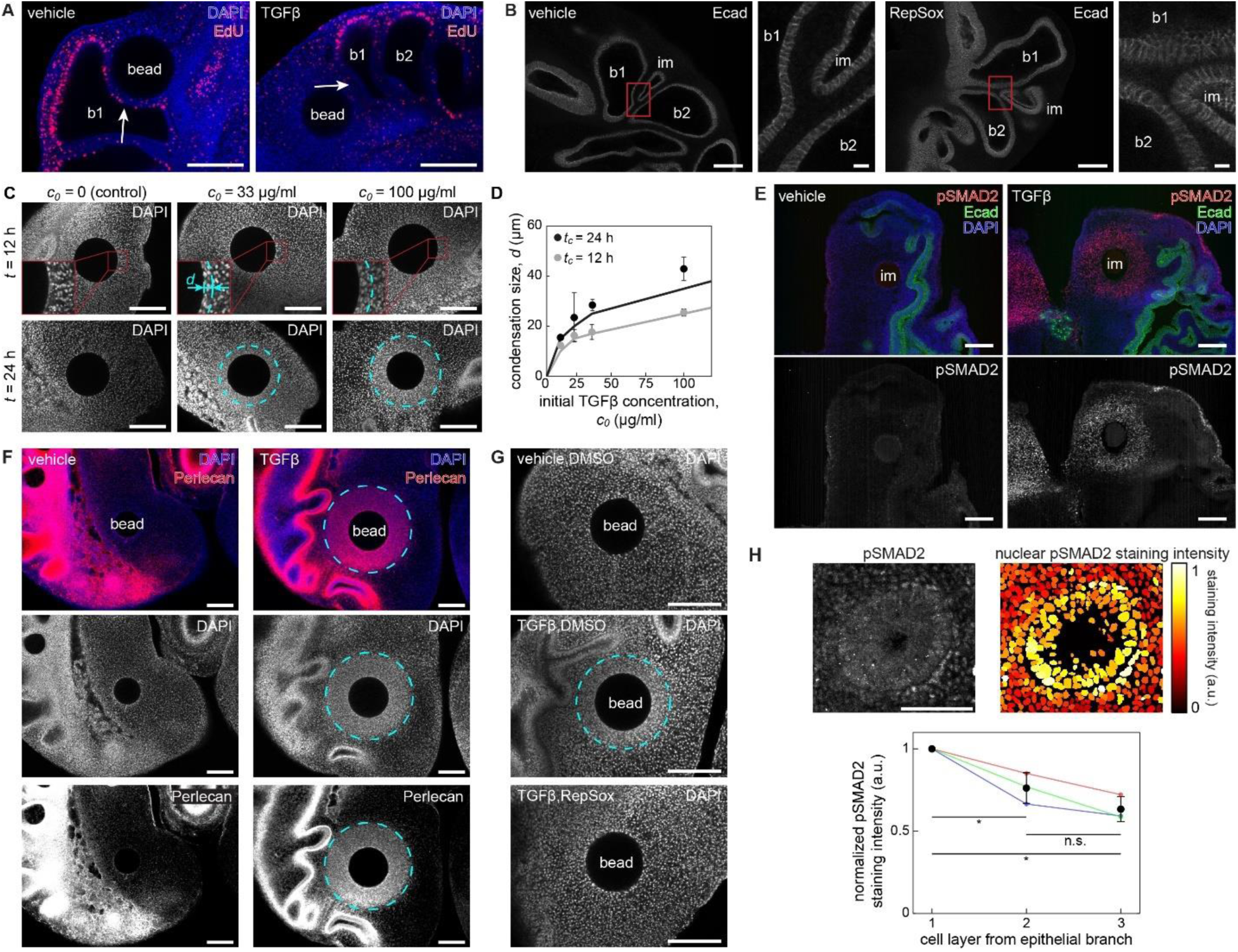
Effects of TGFβ on epithelial and mesenchymal cells in the developing lung – related to Fig. 4. (**A**) Fluorescence images of nuclei (blue) and EdU incorporation (red) in lung explants inserted with TGFβ-containing bead or vehicle control. The beads were inserted proximal to b1. In contrast to control, few EdU-positive mesenchymal cells are observed in TGFβ-containing bead-inserted lung. White arrows denote proximal stalk of b1. (**B**) Fluorescence images of epithelium in branch-implanted lung explants treated with RepSox or vehicle control. The distance between implanted and native branches is small in RepSox-treated lungs. Scale bars, 100 µm and 10 µm (inset). (**C**) Fluorescence images of nuclei in lung explants cultured for 12- or 24-hours after implantation of beads with different TGFβ concentrations. (**D**) Graph showing the sizes of condensations in bead-implanted lung explants with different initial TGFβ concentration and culture time. (**E**) Fluorescence images of nuclei (blue) and pSMAD2 (red) counterstained for E-cadherin (green). Strong circular pSMAD2 staining is observed near TGFβ-containing beads. (**F**) Fluorescence images of nuclei (blue) and perlecan (red) in lung explants inserted with TGFβ-containing bead or vehicle control. Strong perlecan staining is observed in mesenchymal condensations near TGFβ-containing beads. (**G**) Fluorescence images of nuclei in lung explants cultured with or without RepSox after being inserted with TGFβ-containing beads or vehicle control. Mesenchymal condensations are only observed in lungs inserted with TGFβ-containing beads and cultured without RepSox. (**H**) pSMAD2 staining and nuclear-masked pSMAD2 staining intensity near the branch. Scale bar, 50 µm. Subepithelial mesenchymal cells exhibit strong pSMAD2 staining. Cyan dotted lines denote condensations (**D, F, G**). Scale bars, 100 µm (**A, C, E, F, G**).

## Captions for Supplementary Videos

**Supplementary Video S1 – related to Fig. 2**. Live-imaging of cytoplasmic-GFP expressing *E*5 chicken lung. Duration = 20 h. Scale bar, 50 µm.

**Supplementary Video S2 – related to Fig. 5**. Live-imaging of cytoplasmic-GFP expressing *E*6 chicken lung implanted with vehicle-containing bead. Duration = 16 h. Scale bar, 50 µm

**Supplementary Video S3 – related to Fig. 5**. Live-imaging of cytoplasmic-GFP expressing *E*6 chicken lung implanted with TGFβ-containing bead. Duration = 16 h. Scale bar, 50 µm

